# poolHelper: an R package to help in designing Pool-Seq studies

**DOI:** 10.1101/2023.01.20.524733

**Authors:** João Carvalho, Rui Faria, Roger K. Butlin, Vítor C. Sousa

**Author notes:** Corresponding author’s.

## Abstract

Next-generation sequencing of pooled samples (Pool-seq) is an important tool in population genomics and molecular ecology. In Pool-seq, the relative number of reads with an allele reflects the allele frequencies in the sample. However, unequal individual contributions to the pool and sequencing errors can lead to inaccurate allele frequency estimates. When designing Pool-seq studies, researchers need to decide the pool size (number of individuals) and average depth of coverage (sequencing effort). An efficient sampling design should maximize the accuracy of allele frequency estimates while minimizing the sequencing effort. We introduce an R package, *poolHelper*, enabling users to simulate Pool-seq data under different combinations of average depth of coverage, pool sizes and number of pools, accounting for unequal individual contribution and sequencing errors, modelled by parameters that users can modify. *poolHelper* can be used to assess how different combinations of those parameters influence the error of sample allele frequencies and expected heterozygosity. The mean absolute error is computed by comparing the sample allele frequencies obtained based on individual genotypes with the frequency estimates obtained with Pool-seq. Using simulations under a single population model, we illustrate that increasing the depth of coverage does not necessarily lead to more accurate estimates, reinforcing that finding the best Pool-seq study design is not straightforward. Moreover, we show that simulations can be used to identify different combinations of parameters with similarly low mean absolute errors. The *poolHelper* package provides tools for performing simulations with different combinations of parameters before sampling and generating data, allowing users to define sampling schemes that minimize the sequencing effort.

## Introduction

Next Generation Sequencing is an important tool for many biologists, providing access to polymorphism data across a wide range of model and non-model species (Ellegren, 2014). Although the cost of sequencing is continuously decreasing, high coverage sequencing of multiple individuals is still expensive. Furthermore, it is challenging to obtain individual genomic data for certain species (e.g., small organisms) or in certain research areas (e.g., Evolve-and-Resequence experiments). In those instances, next-generation sequencing of pooled samples (Pool-seq) might be the only viable alternative, as it requires less DNA per individual. Additionally, Pool-seq does not require individual tagging of sequences, reducing the laboratory work required for library preparation, while still generating population-level genomic data (Schlötterer, Tobler, Kofler, & Nolte, 2014).

However, Pool-seq introduces new sources of uncertainty in the analysis of genomic data. One common concern is that stochastic variation in amplification efficiency and non-equimolar quantities of DNA in a pool might lead to a loss of accuracy in allele frequency estimations (Anderson, Skaug, & Barshis, 2014; Ellegren, 2014). These differences in DNA concentration, library preparation and amplification efficiency might propagate to cause differences in contribution between pools of individuals when DNA was extracted from multiple batches of individuals and combined into larger pools for library preparation and sequencing (Morales et al., 2019; Ross, Endersby-Harshman, & Hoffmann, 2019). Nevertheless, Pool-seq has been extensively used in a variety of settings (Begun et al., 2007; Ferretti, Ramos-Onsins, & Pérez-Enciso, 2013; Prescott et al., 2015; Zhou et al., 2011). However, there is a lack of tools to simulate and analyse this type of data (but see Kofler, Pandey, and Schlötterer 2011). Two key parameters in the design of a Pool-seq study are the number of individuals in each pool, and the average depth of coverage. These two parameters determine how much the sample allele frequencies are affected by Pool-seq associated errors. On one hand, increasing the number of individuals allows estimating more accurate allele frequencies, but more individuals in the pool might increase the probability of errors associated with unequal individual contribution. On the other hand, increasing the depth of coverage should lead to more reliable estimates, but it can amplify Pool-seq errors and the depth of coverage is usually the limiting resource due to its costs. Simulations of single nucleotide polymorphism (SNP) data accounting for sources of uncertainty with Pool-seq data (e.g., unequal individual contribution) under different sampling schemes can thus provide a tool to help researchers design Pool-seq experiments and to minimize the error associated with the sample allele frequencies.

Here, we introduce an R package (Team, 2020), *poolHelper*, to simulate Pool-seq data according to different sampling designs. Our approach relies on coalescent simulations under neutrality using scrm (Staab, Zhu, Metzler, & Lunter, 2015). The *poolHelper* package provide tools and functions to simulate Pool-seq datasets accounting for potential sources of error, modelled by parameters that users can adjust. This allows comparing the allele frequencies obtained directly from the simulated individual genotypes with the frequencies obtained from Pool-seq data. Since R is a free and collaborative project, users can use available tools to handle, analyse and visualise genomic datasets. Our goal is to provide a flexible method of simulating Pool-seq data, allowing researchers to design their experiments with a better *a priori* knowledge of possible errors associated with Pool-seq, thus contributing to the recognition of Pool-seq as a valuable source of data to reconstruct the evolutionary history of populations.

## Implementation

The main steps of our pipeline follow a relatively simple scheme: coalescent simulations of individual genotypes under a single population model with a constant size, computation of minor-allele frequencies directly from the genotypes, simulation of Pool-seq given the genotypes, and computation of minor-allele frequencies from the Pool-seq data, assuming that it corresponds to the proportion of reads with a given allele. To measure the error associated with Pool-seq we computed the average absolute difference between the actual allele frequencies based on individual genotypes in the sample and the allele frequencies obtained with Pool-seq. Thus, note that we measure the difference between two estimates of the population frequency, one based on the sampled individual genotypes and the other obtained with Pool-seq of the same sample. The *poolHelper* package provides functions to simulate Pool-seq data, under a variety of user-defined conditions. More specifically, users can vary the average coverage, the pool size and the Pool-seq error (see below). Additionally, they can also vary the number of pools used to sequence the population and the sequencing error. By varying all of these conditions, it is possible to assess how they influence the accuracy of allele frequency estimations. No external R objects are needed to use the package. Users can define the mean and variance of the depth of coverage, the number of pools and individuals and the pooling and sequencing errors.

### Coalescent simulations of individual genotypes

To obtain individual genotypes, we used scrm to simulate coalescent gene trees under a model of a single population with constant effective size *N_e_*. To model different effective population sizes and mutation rates, users can vary *θ* = 4*N_e_μ*, where *μ* is the neutral mutation rate per locus per generation. This allows to investigate Pool-seq errors in populations with varying levels of expected genetic diversity, which is proportional to *θ*. We assumed that the sample size was the same for each locus, corresponding to the total number of individuals in the Pool-seq experiment. Additionally, we assumed that the actual haplotypes of all individuals in the pool were known. The effect of pooling is simulated in posterior steps (see next section). To obtain individual genotypes, we assumed random mating in the population and paired haplotypes at each locus at random for each biallelic single nucleotide polymorphic (SNP) site.

### Simulation of Pool-seq data

The steps required to simulate Pool-seq allele frequencies at biallelic SNPs are detailed in Carvalho, Morales, Faria, Butlin, and Sousa (2022). Briefly, we model the depth of coverage at each SNP (i.e., number of reads per site) with a negative binomial distribution (Sampson, Jacobs, Yeager, Chanock, & Chatterjee, 2011), which is defined based on the mean and variance of the depth of coverage across all sites. When simulating the depth of coverage for each site, it is possible to apply a coverage-based filter, removing all sites with coverage below and above two user-defined thresholds. At each retained SNP, we modelled heterogeneity in the contribution of each individual to the DNA pool, by assuming that the number of reads from each individual follows a multinomial distribution, with the expected proportion of reads from each individual following a Dirichlet distribution. To account for situations where DNA extraction was performed for several batches of individuals and the extracted DNA from each batch merged into a single pool, we also model the possibility of uneven contributions between pools. This was also assumed to follow a multinomial distribution, with the expected proportion of reads from each pool (i.e., batch) following a Dirichlet distribution. In both instances, the dispersion of the Dirichlet distribution is controlled by a Pool-seq error parameter, following Gautier et al. (2013). This parameter affects the variance of the proportion of reads from each individual, which captures the random unequal individual contribution to the pool (Gautier et al., 2013). Larger Pool-seq errors lead to larger variance, resulting in more unequal contributions from individuals and pools. Although the selection of an appropriate Pool-seq error might be potentially hard, given its unknown nature, we previously estimated values ranging from 24 to 236, with posterior means for two different models of 182 and 102 (Carvalho et al., 2022). Thus, the Pool-seq errors used here are within the reasonable ranges for this parameter (see Figure 1 for an example of how different Pool-seq errors impact individual contribution). Note that this model assumes that all individuals are expected to contribute the same number of reads, with errors due to unequal contribution modelled through dispersion parameters that affect the variance. Finally, to model the effect of putative sequencing and mapping errors, we considered that the allele in the reads of the pool could differ from the actual genotype. Thus, for each individual, the number of reads with the alternative allele follows a binomial distribution, assuming that with a given error rate, a reference allele will incorrectly be called an alternative allele or vice-versa. A commonly used filter can also be applied, discarding SNPs with less than the required number of minor-allele reads. The allele frequencies estimated for the Pool-seq data correspond to the proportion of reads with the alternative allele.

**Figure 1:**
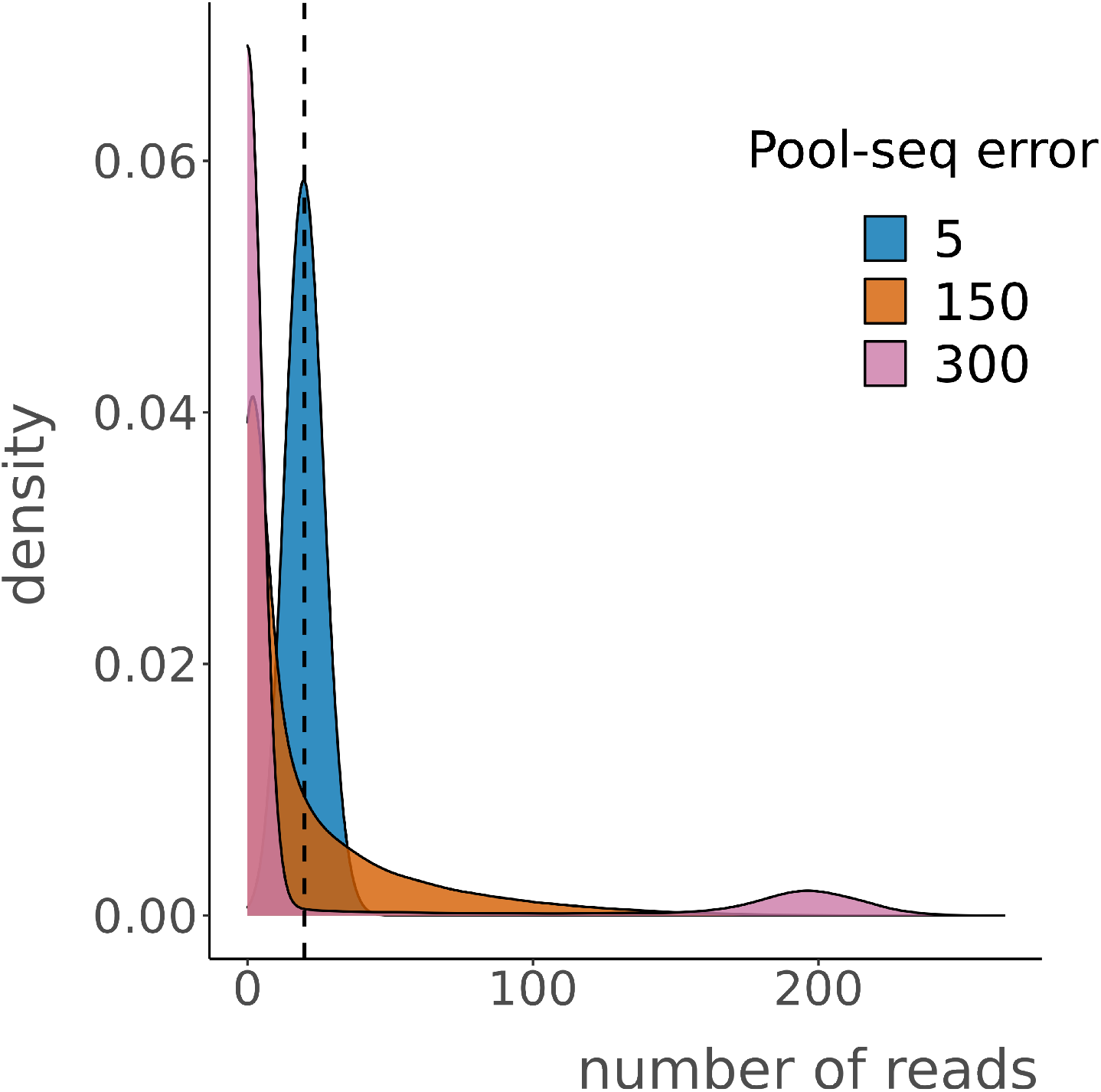
Impact of Pool-seq errors in individual contribution. We simulated a situation where a single pool of 10 individuals was sequenced at 200x coverage. The number of reads contributed by each individual was simulated under three different Pool-seq errors: 5, 150 and 300. The dashed line indicates the expected contribution if all individuals contributed equally. Note that increasing Pool-seq errors lead to deviations from the expected value and a marked increase in the number of individuals contributing zero (or close to zero) reads. Note also that with a high Pool-seq error (300) there are some individuals contributing ~200 reads.

### Measuring error of estimates

To measure the error of Pool-seq estimates of allele frequencies or expected heterozygosity, we compared the estimates obtained from the individual genotypes in the sample with the estimates obtained from Pool-seq. We calculate the mean absolute error as:

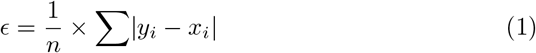

where *n* indicates the total number of SNPs. When calculating the error of Pool-seq estimates of allele frequencies, *x_i_* and *y_i_* correspond to the frequencies of the alternative allele at the *i^th^* SNP in the sample, obtained with individual genotypes (*x_i_*) or with Pool-seq (*y_i_*). When measuring the error of expected heterozygosity, *x_i_* and *y_i_* represent the expected heterozygosity obtained based on the sample of either individual genotypes (*x_i_*) or Pool-seq (*y_i_*).

### Main functionality

The *poolHelper* package allows users to compute the mean absolute error of allele frequencies and expected heterozygosity under a variety of conditions. Users can vary the mean depth of coverage and the associated variance, the value of the Pool-seq error and the number of sampled individuals. Additionally, it is possible to evaluate the effect of combinations of parameters, for instance, various mean depths of coverage combined with several Pool-seq error values. Thus, the *poolHelper* package provides users with a tool to aid in the design of pooled sequencing experiments, by allowing researchers to evaluate the best strategy, in terms of pool sizes or depth of coverage, to obtain accurate estimates of allelic frequencies, while minimizing the sampling effort and costs.

#### Effect of sequencing a single or multiple pools

One critical consideration is whether DNA extraction should be performed on multiple batches of individuals, combining several of them into larger pools for library preparation and sequencing, or on a single batch of individuals. Users can test the effect of using multiple or a single batch of individuals by using the ”maePool” function. This function computes the mean absolute error for a given sample size sequenced using a single or multiple pools (Figure 2). By varying the mean coverage and the Pool-seq error, it is possible to evaluate the effect of using a single or multiple pools under different conditions.

**Figure 2:**
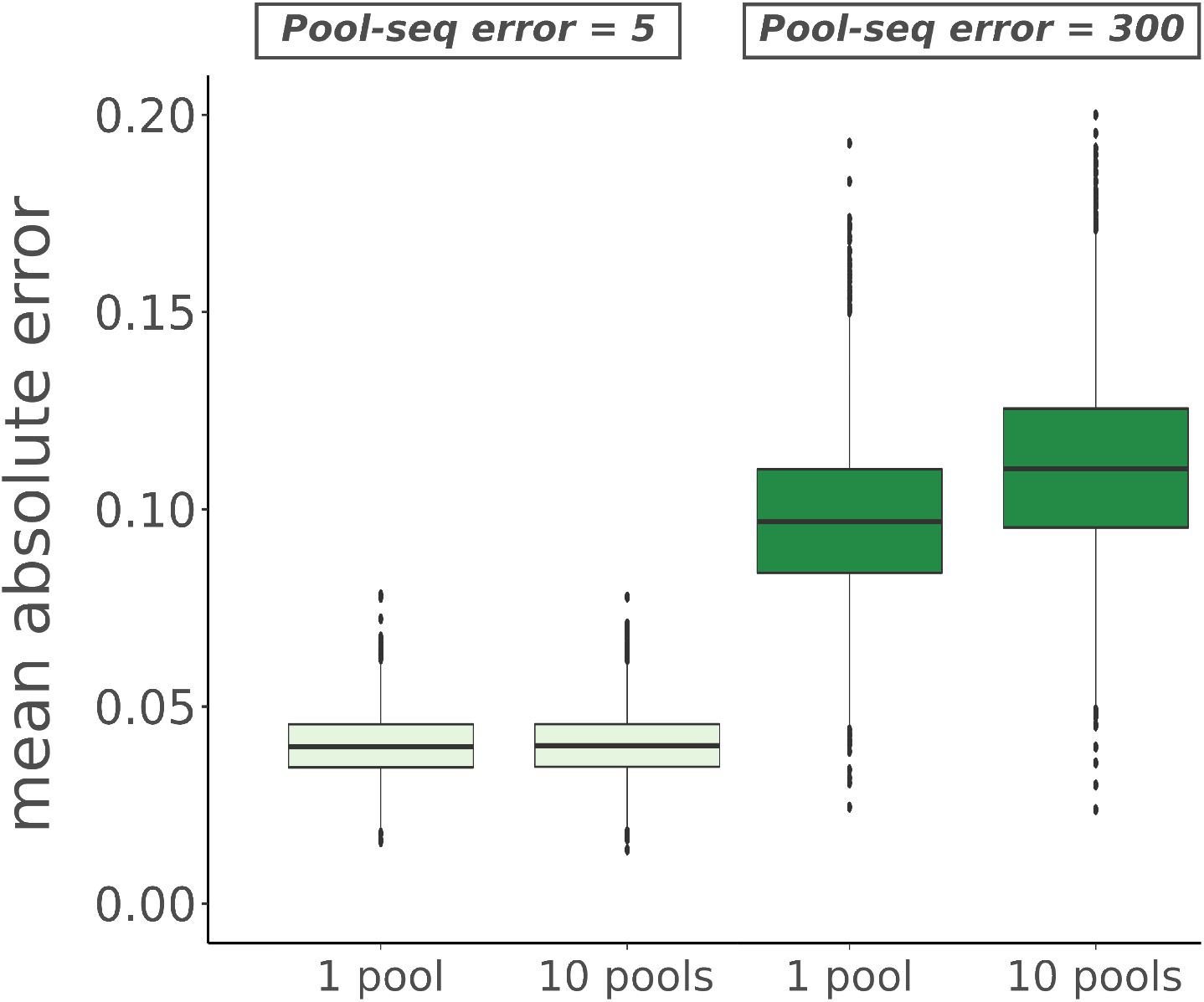
Mean absolute error between the allele frequencies obtained from the individual genotypes in the sample and those obtained from Pool-seq data using a single or multiple pools. Pool-seq data was simulated for 50 individuals, sequenced with average coverage of 50x using a single pool or 10 pools, each with 5 individuals and assuming a low Pool-seq error (5) and a high Pool-seq error (300). The same Pool-seq error value was used to model the dispersion among pools and individuals. The y-axis represents the mean absolute error between the allele frequencies estimates and the x-axis indicates the number of pools used to sequence the sample.

#### Impact of mean depth of coverage

Another critical decision is defining the mean depth of coverage used to sequence a pool of individuals. The ”maeFreqs” function implements the calculation of the mean absolute error between allele frequencies computed from genotypes and Pool-seq allele frequencies simulated under different mean depth of coverage. By varying the mean depth of coverage and the associated variance, users can determine which coverage produces more accurate allele frequency estimates for a given sample size and Pool-seq error (Figure 3).

**Figure 3:**
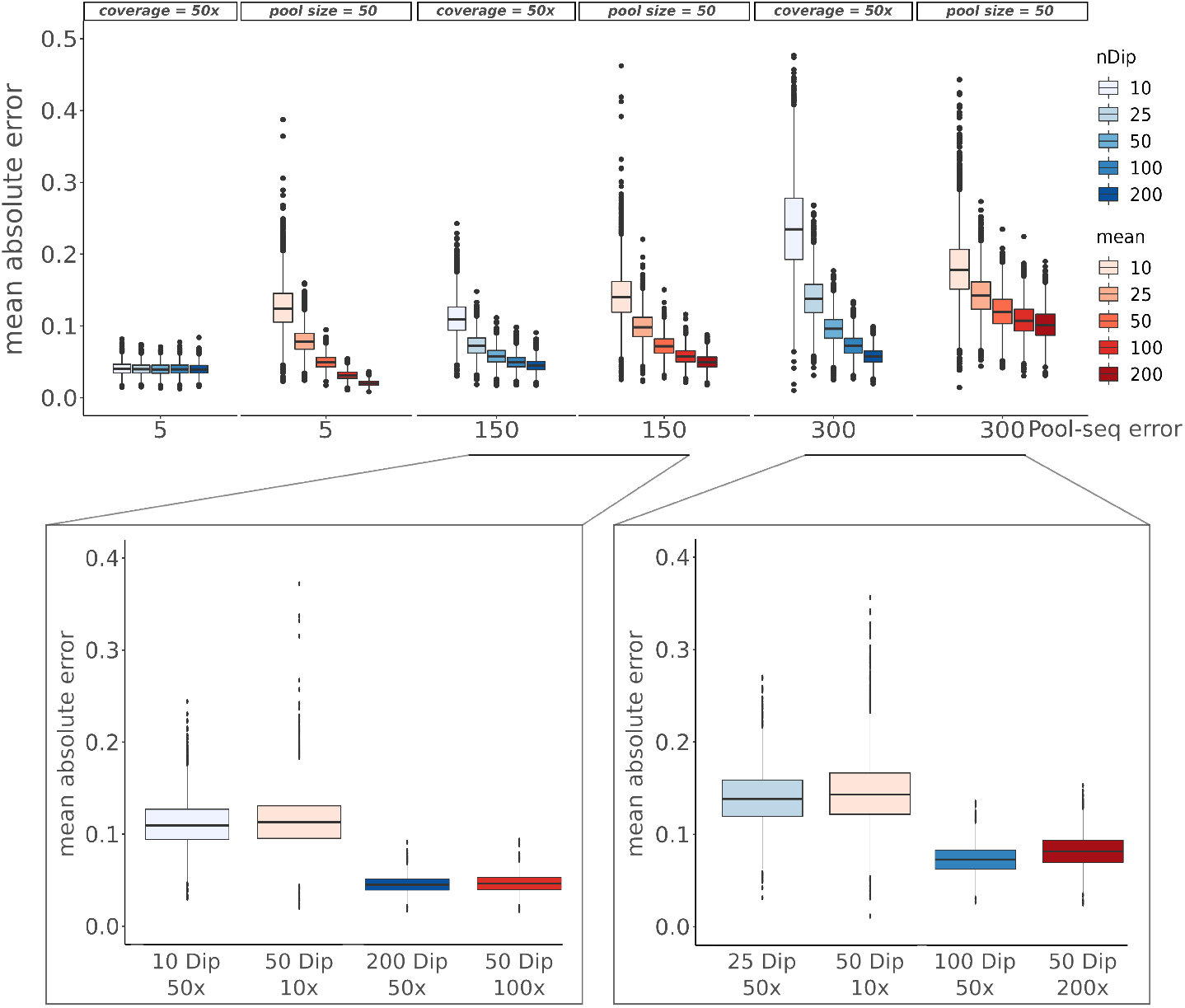
Mean absolute error between the allele frequencies obtained from the individual genotypes in the sample and those obtained from Pool-seq data under a variety of conditions. For all conditions, sites with less than two minorallele reads were removed. In all plots, the y-axis represents the mean absolute error between the allele frequencies estimates. The top panel shows the mean absolute error for three different Pool-seq error values (x-axis). For each plot, either the pool size or the coverage were fixed (the fixed value is indicated on the top of each plot). Thus, when pool size was fixed, the average coverage varied and vice-versa. In the bottom panel, we highlight comparisons that lead to similar mean absolute errors for intermediate values of Pool-seq error (150 in the bottom left panel) and high Pool-seq error (300 in the bottom right panel). In all plots, the pool size, defined by the *nDip* parameter, is represented in shades of blue, with darker shades indicating a bigger pool and the average coverage, defined by the mean parameter, is represented in shades of red, with darker shades indicating higher coverage.

#### Impact of pool sizes

When designing a Pool-seq experiment, it is essential to define the number of individuals to include in the pool, i.e., the pool size. The calculation of the mean absolute error between allele frequencies for different pools sizes can be carried out using the ”maeFreqs” function. This allows users to evaluate what is the optimal pool size for a fixed coverage and/or Pool-seq error (Figure 3). Thus, the ”maeFreqs” function allows users to decide how many individuals to pool to obtain the most accurate allele frequencies estimates for a given mean depth of coverage.

#### Example of an effective Pool-seq design using simulations

By performing simulations in a single panmitic population, assuming that Pool-seq error is intermediate to high (150 or 300) and after applying a commonly used filter (removing sites with less than two minor-allele reads), it is not obvious that one should always increase the average depth of coverage per individual in the pool (Figure 3). For instance, when Pool-seq error is 150, we observe the same mean absolute error with a pool of 50 individuals sequenced at 10x than with a pool of 10 individuals sequenced at 50x. This suggests that it may be more cost-effective to use a pool of 50 individuals at 10x (expected individual contribution of 10/50) than using fewer individuals with a higher expected coverage per individual. This holds true for larger pool sizes and depths of coverage, given that we also get the same mean absolute error when comparing a pool of 200 individuals sequenced at 50x with a pool of 50 individuals sequenced at 100x (Figure 3). If Pool-seq error is even higher (i.e., 300) a pool of 100 individuals sequenced at 100x leads to a slightly lower mean absolute error than a pool of 50 individuals sequenced at double the coverage (200x) (Figure 3). Thus, similar errors of allele frequencies in the sample can be obtained with different combinations of pool sizes and average depth of coverage. Therefore, the design of an effective Pool-seq study is not straightforward and an a priori simulation study can help assess an efficient sampling scheme to obtain accurate allele frequencies while minimizing the sequencing effort (mean depth of coverage).

## Conclusions

We present an R package, *poolHelper*, to simulate pooled sequencing data under a model of a single panmitic population, and compute the error in sample allele frequencies and expected heterozygosity obtained with Pool-seq for different study designs and commonly used filters (e.g., filters on minimum and maximum depth of coverage and on minimum number of minor-allele reads). The package relies on coalescent simulations performed with *scrm* (Staab et al., 2015). Currently, data is simulated under a single population with a constant effective population size. However, users can modify *poolHelper* functions to simulate data under more complex demographic history models. This package is implemented in the R environment, providing tools for data visualisation, allowing users to produce graphics and quickly visualize the effect of multiple combinations of Pool-seq related parameters. The *poolHelper* package’s vignette contains a comprehensive explanation of the functions included in this package, as well as examples detailing its usage.

## Author Contributions

J.C. and V.C.S. conceptualized and designed poolHelper; J.C. implemented the package in R with supervision and input of V.C.S; R.K.B and R.F. discussed preliminary results and provided suggestions that improved the functionality of the package; J.C. wrote the first draft, with comments from all authors. All authors contributed critically to the drafts and gave final approval for publication.

## Acknowledgements

We thank Beatriz Portinha for suggesting the package name. This work was funded by the strategic project UIDB/00329/2020 granted to cE3c from the Portuguese National Science Foundation — Fundação para a Ciência e a Tecnologia (FCT). JC was supported by an FCT Ph.D. scholarship (PD/BD/128350/2017). RF is funded by a FCT CEEC (Fundação para a Ciênca e a Tecnologia, Concurso Estímulo ao Emprego Cientfíco) contract (2020.00275.CEECIND) and by a FCT research project (PTDC/BIA-EVL/1614/2021). RKB was funded by the European Research Council (ERC-2015-AdG-693030-BARRIERS). VCS was supported by Human Frontier Science Program (RGY0081/2020) and by FCT (CEECINST/00032/2018/CP1523/CT0008). We thank the National Network for Advanced Computing (RNCA) and INCD (https://incd.pt/) for access and use of the computing infrastructure, funded by FCT to VCS (2021.09795.CPCA).

## Data Availability

The package poolHelper source code is hosted at https://github.com/joao-mcarvalho/poolHelper, along with the package tutorials and vignettes. The source code has been archived with Zenodo at https://doi.org/10.5281/zenodo.7520303. This will be released as an R package in the CRAN repository. There is no other data associated with this paper.

